# Differential Host Gene Signatures in Response to *Mycobacterium tuberculosis* Infection

**DOI:** 10.1101/2020.02.19.955203

**Authors:** Loubna Tazi, Ping Wang, Myriam Fornage

**Affiliations:** Department of Medical Microbiology & Immunology, School of Medicine, University of California Davis, Davis, CA 95616; Brown Foundation Institute of Molecular Medicine, McGovern Medical School, The University of Texas Health Science Center at Houston, Houston, TX 77030

## Abstract

Tuberculosis (TB) represents a global public health threat and is the leading cause of morbidity and mortality worldwide. Effective control of TB is complicated with the emergence of multidrug resistance. Yet, there is still a fundamental gap in understanding the complex and dynamic interactions between different *Mycobacterium tuberculosis* strains and the host. In this project, we investigated the host immune response to different *M. tuberculosis* strains, including avirulent or virulent and rifampin-resistant or isoniazid-resistant strains in THP-1 cells. We identified major differences in the gene response profiles in response to infection with these strains. The expression of IDO1 and IL-1β in the infected cells was stronger in all virulent *M. tuberculosis* strains. The most striking result was the overexpression of many interferon-stimulated genes (ISGs) in cells infected with the isoniazid-resistant strain, compared to the rifampin-drug resistant strain and the drug-sensitive strain. A transcription regulation analysis of the differentially expressed genes in infected THP-1 cells implicated two major transcription factors, NF-κB and STAT1. The differentially expressed ISGs in response to the isoniazid-resistant *M. tuberculosis* strain were associated with STAT1 signaling, while the expression of many cytokines, such IL-1β, was associated with NF-κB signaling. Our data suggest that the isoniazid-resistant *M. tuberculosis* strain preferentially activates STAT1 in response to cGAS-STING activation and induces a host immune response signature that is characteristic of isoniazid resistance. This study has a potential to provide important new insights into TB pathogenesis and to characterize host gene signatures specifically involved in isoniazid-resistant TB.

## INTRODUCTION

Tuberculosis (TB) represents a disease of global public health importance and a leading cause of morbidity and mortality worldwide, surpassing HIV/AIDS (1, 2). One third of the world’s population is currently infected with *Mycobacterium tuberculosis*, with more than 10 million new cases of active TB reported worldwide resulting in more than 1.6 million deaths annually. The emergence of drug-resistant TB and the increasing incidence of multidrug-resistant (MDR) and, more recently, of extensively drug-resistant (XDR) and totally drug-resistant (TDR) *M. tuberculosis* strains are a highly significant public health threat and jeopardize current efforts to TB control and prevention (3-5).

Besides the ability to acquire mutations in target genes conferring resistance to antimicrobial drugs, *M. tuberculosis* strains also show a high degree of intrinsic resistance to most common antibiotics (6). This allows the mycobacteria to efficiently resist anti-tuberculous treatment. As a consequence, the molecular characterization of drug-resistant *M. tuberculosis* strains remains technically challenging (7-9). There is an urgent need to develop alternative and more sensitive approaches to identify *M. tuberculosis* strains in order to implement better measures to minimize the acquisition of further drug resistance, prevent TB transmission, and ultimately improve TB control and prevention strategies.

The *M. tuberculosis* cell wall has a complex composition and structure. It is considered to be a major virulence factor and to promote the natural resistance of *M. tuberculosis* to antibiotics (10). Mycolic acids represent the hallmark component of the *M. tuberculosis* cell wall and their biosynthesis and regulation are the targets of isoniazid (INH), one of the cornerstone drug of TB treatment (11, 12). Interestingly, the cell wall is significantly thicker in drug-resistant *M. tuberculosis* strains compared to drug-sensitive strains, further underscoring the importance of mycolic acids biosynthesis in INH-resistant *M. tuberculosis* strains (13).

Cell wall lipids in *M. tuberculosis* play a significant role in modulating the host immune response, as well as its pathogenic processes and virulence (14-17). However, our understanding of the host-*M. tuberculosis* interaction is still limited, and many fundamental gaps remain in how this interaction could be altered in response to different *M. tuberculosis* strains. Infections with different *M. tuberculosis* genotypes have major impacts on host-pathogen interaction, which can lead to substantial differences in the host immune responses (18-20). Some mycobacterial lineages (e.g., Beijing) even showed polymorphic properties in immune response genes, suggesting a possible human-pathogen co-evolution (21). The immune response of infected macrophages also showed a wide variation in the response to different *M. tuberculosis* lineages (22). However, further work is needed to explore the host immune response to different *M. tuberculosis* strains. In this study, we tackled this question and investigated the host immune response to different *M. tuberculosis* strains, including avirulent or virulent and rifampin-resistant or isoniazid-resistant strains in THP-1 cells. Shifting the focus on the host-pathogen interaction in *M. tuberculosis* infection and the identification of specific host responses to *M. tuberculosis* strains have the potential to develop biomarkers for novel and more sensitive diagnostic tools for TB. The identification of TB patients at an early stage, particularly those carrying drug-resistant strains, is crucial in order to begin an appropriate therapy as quickly as possible and improve our efforts for TB control.

## MATERIALS AND METHODS

### Cell line treatment and *M. tuberculosis* infection

For this study, we used human monocytic THP-1 cells (American Type Culture Collection; Catalog no. TIB-202). The cells were maintained in RPMI 1640 medium supplemented with 10% heat-inactivated fetal bovine serum (FBS) at 37°C and 5% CO_2_ in a humidified incubator. THP-1 cells were subsequently differentiated for 48 hours with 10 ng/ml phorbol 12-myristate 13-acetate (PMA). The cells were then infected with four different reference mycobacterial strains (H37Ra, H37Rv, H37Rv-INH-R, H37Rv-RIF-R), kindly provided by Dr. Lisa Y. Armitige (Heartland National TB Center). Both H37Ra (American Type Culture Collection; Product no. 25177) and H37Rv (American Type Culture Collection; Product no. 27294) are drug-sensitive, whereas H37Rv-INH-R (American Type Culture Collection; Product no. 35822) and H37Rv-RIF-R (American Type Culture Collection; Product no. 35838), both of which are derivatives of H37Rv, are resistant to isoniazid and rifampin, respectively. We used two time points for infection (4 and 24 hours) in order to monitor differences in gene expression patterns for differentially expressed genes. In addition to the infected THP-1 samples, we also used uninfected control cells (mock-infected with PBS) as negative controls.

### RNA isolation and processing

Total RNA was extracted from controls and infected cells (10 samples in total, with 5 samples for each time point), using Trizol according to the manufacturer’s protocol. RNA was then suspended in RNase-free water and stored at −80°C until further use. RNA quality was assessed to verify its integrity using Agilent 2100 Bioanalyzer (Agilent Technologies) and RNA quantity was evaluated by spectrophotometry using NanoDrop 2000 (NanoDrop Technologies). All RNA samples showed good RNA yield and no RNA degradation. Total RNA was then reverse transcribed to cDNA, amplified, labeled and hybridized to separate arrays using the GeneChip Human Exon 1.0 ST Array (Affymetrix, USA), according to the manufacturer’s instructions. For transcriptional profiling, we used triplicate hybridization assays in our microarray experiments in order to assess variability among independent labeling reactions and hybridizations. Quality control of the hybridized arrays was also performed for each sample. A visual inspection of the scanned images was conducted looking for any defects, areas of high background, or areas of low signal. The spike-in controls were checked as well to examine for hybridization uniformity.

### Microarray data analysis

Data from all 10 samples were preprocessed, summarized at the transcript-cluster (gene) level, and RMA normalized using Affymetrix Power Tools. Prior to differential expression analysis, low-variability genes were filtered out, leaving 13,460 genes. Differential expression analyses were conducted using the R/Bioconductor package *limma* (23). Significant differential expression was defined by an adjusted p-value of less than 0.05. The visualization of the identified differentially expressed genes in the volcano plots was performed using base R graphics.

## RESULTS

In this study, we generated gene expression profiles of THP-1 cell lines infected with different reference *M. tuberculosis* strains (H37Ra, H37Rv, H37Rv-INH-R, H37Rv-RIF-R). The choice for this study design allowed us to focus on *M. tuberculosis* strain-specific alterations in the transcriptional response and to avoid putative confounding differences that might be related to the genetic variability of clinical *M. tuberculosis* strains and macrophages isolated from different subjects. The multidimensional scaling (MDS) plot for the microarray expression data clearly showed well-separated grouping according to infections vs mock infections and time points of infections, and also that replicate samples clustered together (Fig. 1).

**Figure 1:**
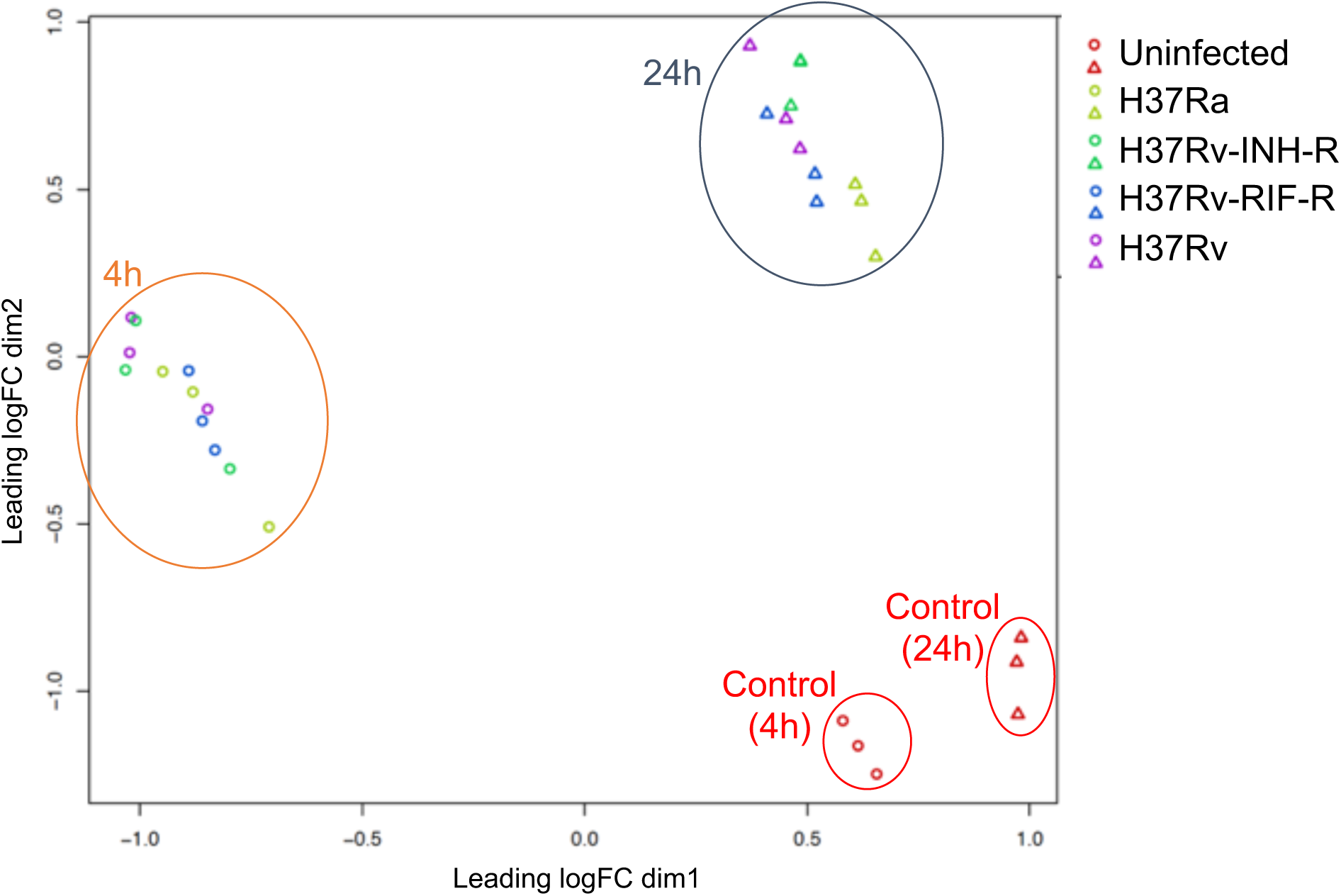
MDS Plot for differential gene expression of *M. tuberculosis*-infected vs mock-infected THP-1 cell lines at 4 h and 24 h post-infection.

Among the total number of differentially expressed genes between infected THP-1 cells and mock-infected cells, we found major differences in those genes, depending on the *M. tuberculosis* strain used for infection. A higher number of differentially expressed genes was detected in infected cells with all virulent *M. tuberculosis* strains (H37Rv, H37Rv-INH-R, H37Rv-RIF-R), compared to the infection with the non-virulent strain (H37Ra) (Fig. 2). This difference was even more pronounced at 24 h post-infection when we performed pairwise comparisons in infected cells (Fig. 3). Based on this finding, the subsequent analyses were therefore based on that time of infection (24 h), instead of 4 h post-infection. A total of 3,622 differentially expressed genes (p<0.05) was identified in response to H37Rv-INH-R infection.

**Figure 2:**
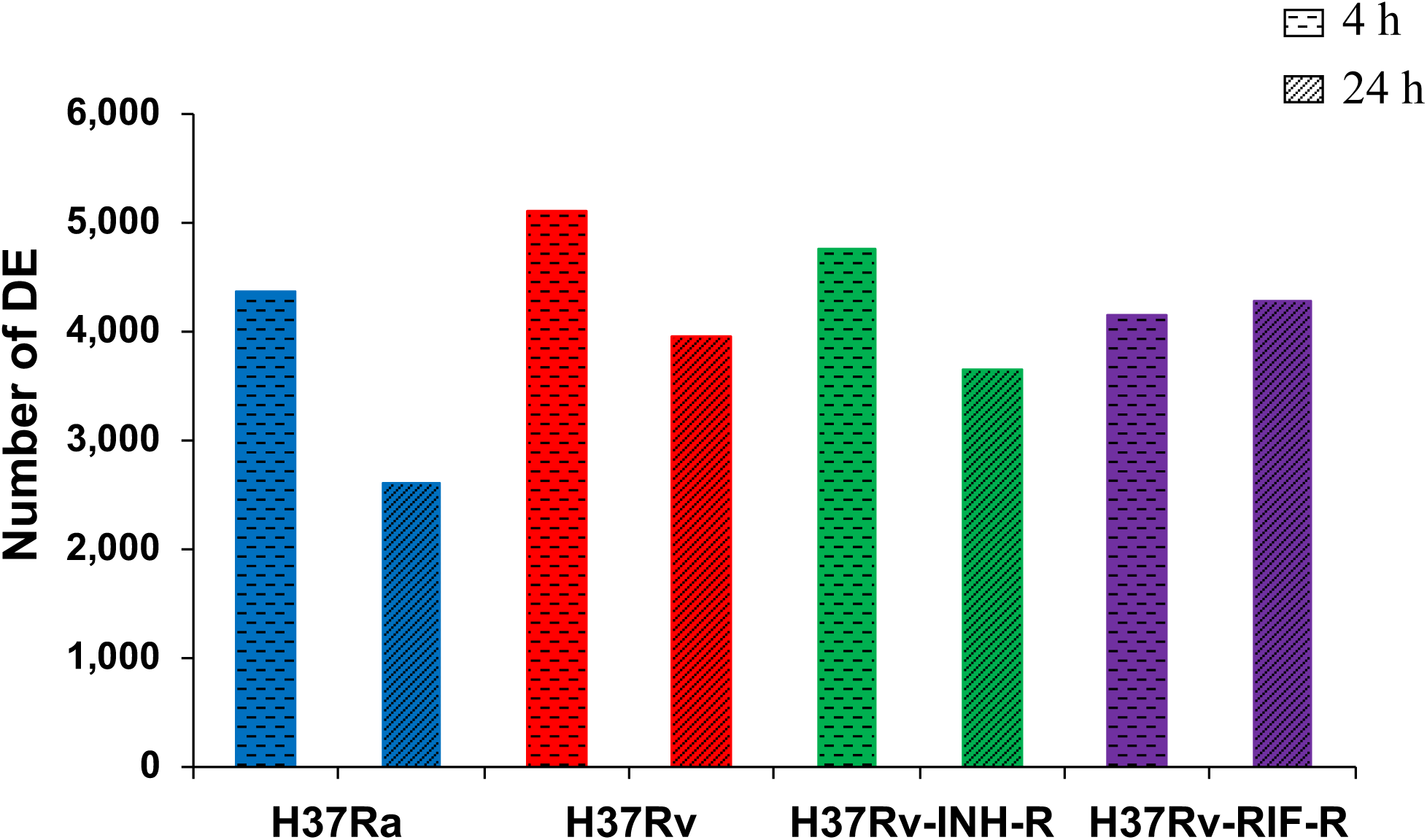
Number of differentially expressed genes in *M. tuberculosis*-infected THP-1 cell lines, relative to mock-infected cells, at 4 h and 24 h post-infection (p<0.05).

**Figure 3:**
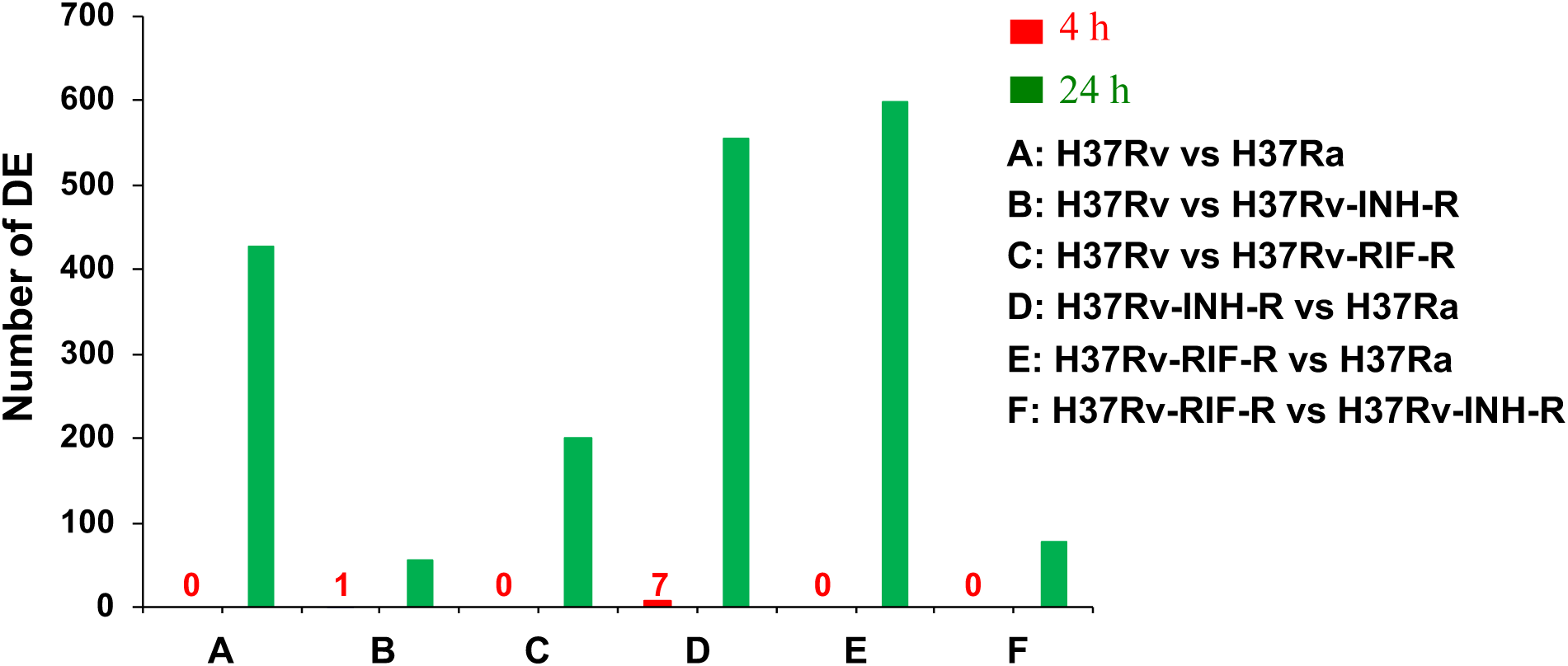
Number of differentially expressed genes in the pairwise comparisons of *M. tuberculosis*-infected THP-1 cell lines at 4 h and 24 h post-infection (p<0.05).

At 24 h post-infection, we identified differences in the gene response profiles between the strains. The expression of IDO1 and IL-1β was enhanced in all infections with the virulent reference strains, except H37Ra (Fig. 4). However, the most striking and unexpected result was the overexpression of many interferon-stimulated genes (ISGs) in the THP-1 cells infected with H37Rv-INH-R, in comparison to cells infected with the other *M. tuberculosis* strains (Fig. 5). Our data also showed that protein kinase R (PKR/eIF2aK2) is most strongly induced in THP-1 cells infected with H37Rv-INH-R. A subset of the differentially expressed host genes is shown in figure 5. In this figure, we can see a panel of differences in the host expression profile. The gene expression changes between infections with H37Rv-INH-R vs the other strains are also shown in the volcano plots (Fig. 6).

**Figure 4:**
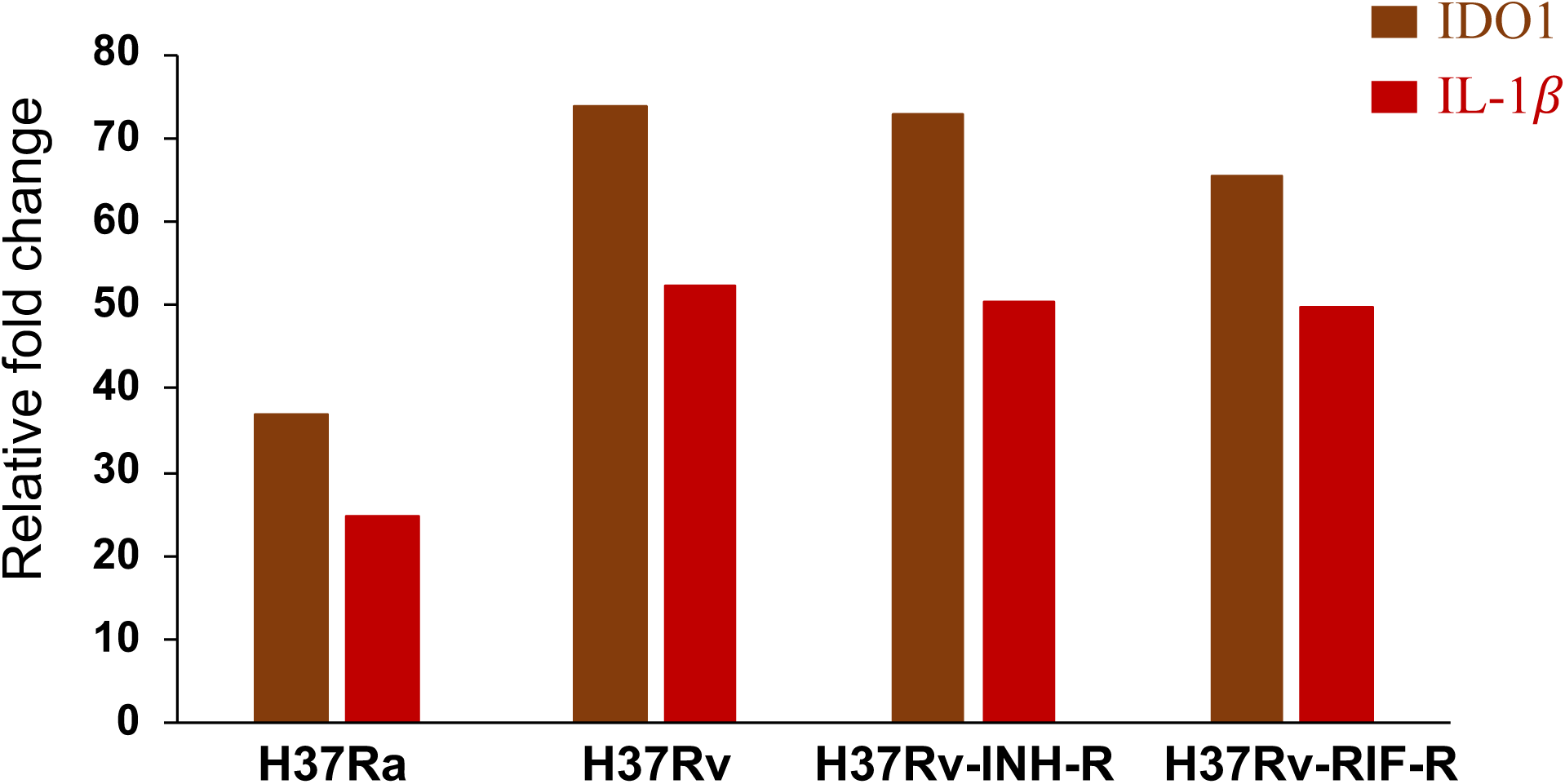
Fold change in the expression of IDO1 and IL-1β in THP-1 cell lines infected with reference *M. tuberculosis* strains (24 h post-infection; p<0.05).

**Figure 5:**
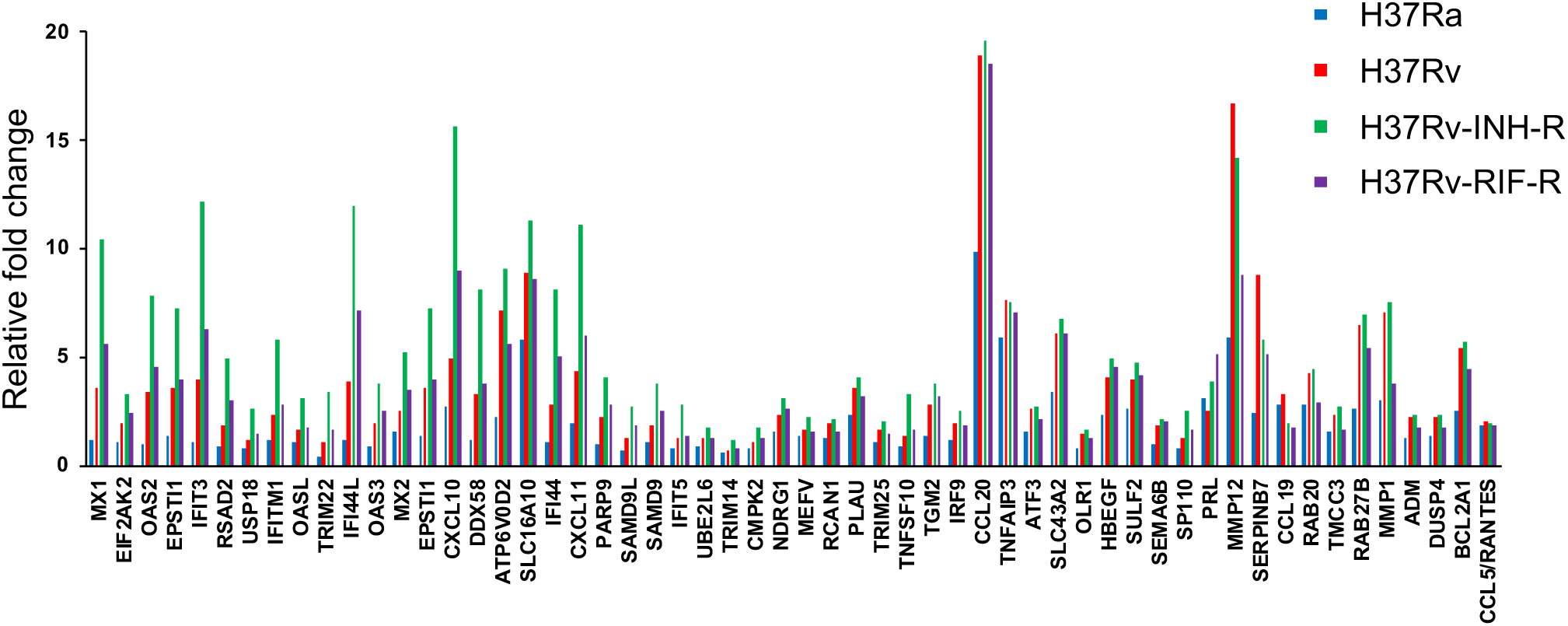
Fold change in gene expression in THP-1 cell lines infected with reference *M. tuberculosis* strains (24 h post-infection; p<0.05).

**Figure 6:**
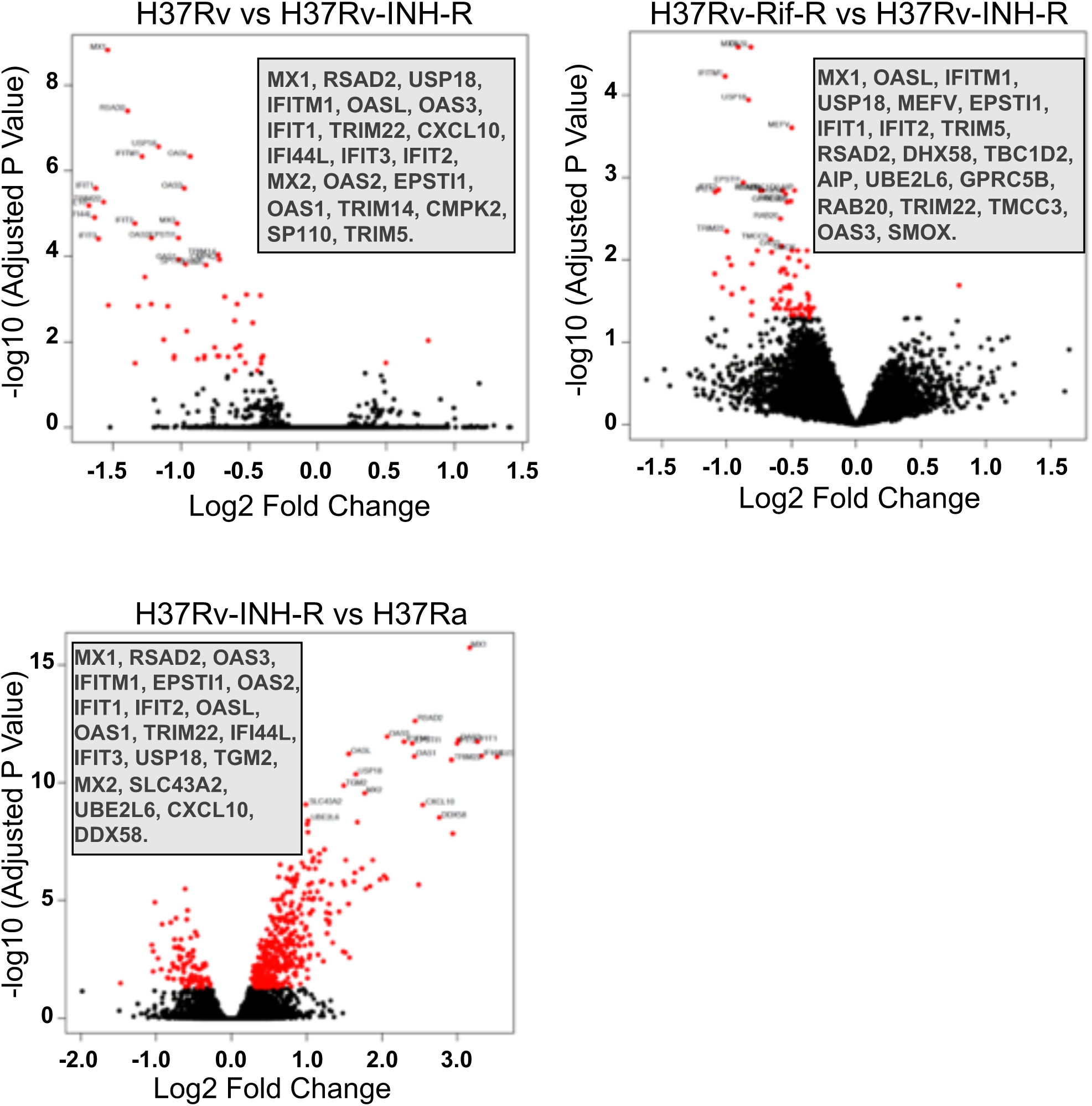
Volcano plots displaying differential expression genes in THP-1 cells in three different comparison groups (24 h post-infection; p<0.05). The 20 highly significant differentially expressed genes in each plot are indicated in the insets.

To determine the key transcription factors that potentially regulate the differentially expressed genes in infected THP-1 cells, we performed a transcription regulation analysis using pathway relation network (pathway-net: http://pathwaynet.princeton.edu/) (24). Two major transcription factors, nuclear factor kappa-light-chain-enhancer of activated B cells (NF-κB) and signal transducer and activator of transcription 1 (STAT1), were identified in this analysis. Our analysis showed that the differentially expressed ISGs in H37Rv-INH-R were associated with STAT1 signaling, while the expression of many cytokines, such IL-1β, was associated with NF-κB signaling.

## DISCUSSION

Our study showed major differences in the host response to different *M. tuberculosis* strains. We also showed that the infection with H37Rv-INH-R induced an unexpected overexpression of ISGs in THP-1 cells. Our data indicated that STAT1 is the major transcription factor associated with the differentially expressed genes (e.g., ISGs) in response to H37Rv-INH-R infection.

Multiple recognition and signaling pathways for the host innate immune responses to *M. tuberculosis* have been documented and characterized (25). The main signaling pathways in *M. tuberculosis* are TLR2-MyD88 and cGAS-STING. In this study, the overexpression of ISGs in the cells infected with H37Rv-INH-R suggests a stronger induction of STAT1 as a result of cGAS-STING activation, rather than TLR2-MyD88 activation in the differential host response to INH-resistant TB. In addition to the overexpression of ISGs in the cells infected with the isoniazid-resistant *M. tuberculosis* strain, we also found a higher induction of PKR/eIF2aK2 in those cells. This result is even more exciting, since there has been recently an increasing interest in PKR/eIF2aK2, a key player in the innate immune response to viral infections, for its potential ability to improve TB control (26-29). Recent studies have also shown an important role of NF-κB dynamics in TB pathogenesis (30, 31). NF-κB was the other major transcription factor associated with the differentially expressed genes, which was identified in this study.

Resistance to INH is linked to all multidrug-resistant forms of TB (MDR, XDR, TDR). Several studies have shown that INH resistance is acquired first, followed by resistance to rifampin and the other anti-TB drugs (32-36), which highlights the significant impact of INH in the success of treatment in active and latent TB and warrants further efforts to understand the global burden of INH-resistant TB (37). Fig. 7 shows our proposed model for the host signaling response to the different reference *M. tuberculosis* strains (H37Ra, H37Rv, H37Rv-INH-R, H37Rv-RIF-R).

**Figure 7:**
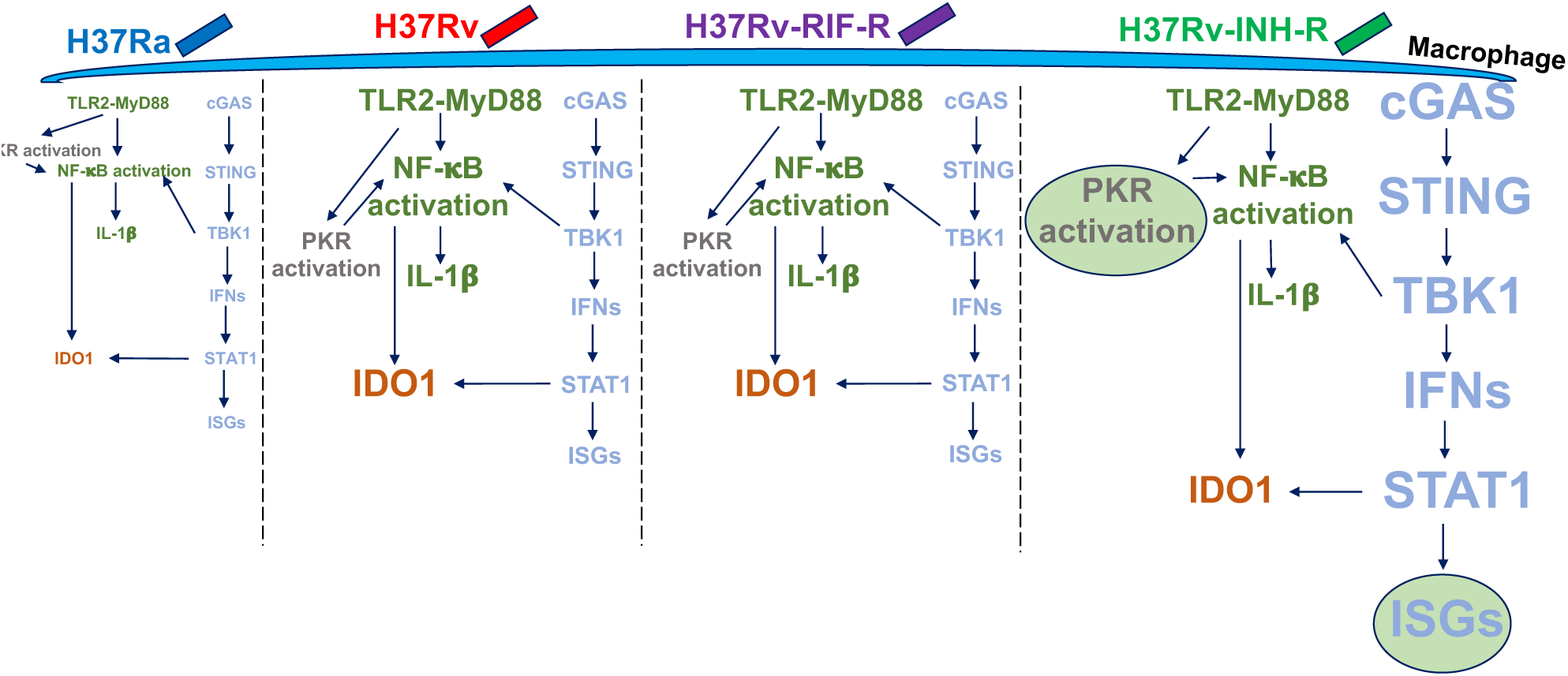
Proposed model for host signaling response to reference *M. tuberculosis* strains. H37Rv-INH-R induces the highest expression of ISGs and PKR, compared to the other strains, as indicated by the circles and the big font size.

The mechanism of INH resistance is complex and has been thoroughly studied in an attempt to improve early diagnosis of INH-resistant *M. tubeculosis* strains (36, 38-44). It is mainly mediated by mutations in the *katG* gene or in the *inhA* regulatory regions (45). INH is activated by the catalase-peroxidase encoded by *katG*. This process interferes with the biosynthesis of mycolic acids by inhibiting NADH-dependent enoyl-ACP reductase encoded by *inhA*. Mutations in other genes have been also associated with INH resistance, however they are not as common as the ones in *katG* (42 to 95%) and *inhA* (6 to 43%) in *M. tuberculosis* clinical strains and their mode of action in INH resistance has not been fully elucidated (46). Nevertheless, not all INH-resistant strains harbor defined genetic mutations associated with resistance to this drug, which complicates efforts to identify those strains. There are also INH resistance-conferring mutations that cannot be detected by current molecular diagnostics approaches (33). In addition, some studies indicated that some *M. tuberculosis* strains acquire drug resistance at higher rates, suggesting a higher mutation rate in these strains and a higher probability that these strains will develop multidrug resistance (47, 48).

Given our results, the differential host immune response to different *M. tuberculosis* strains (e.g. overexpression of ISGs in the cells infected with H37Rv-INH-R) provides opportunities to develop host biomarkers for INH resistance and defines a new facet of host-pathogen interactions that differentially regulate signaling pathways in response to INH-resistant and other *M. tuberculosis* strains. The characterization of such host response signatures has the potential to develop novel diagnostic tools and improve our understanding of the complex and dynamic interactions between *M. tuberculosis* strains and the host.

## ACKNOWLEDGMENTS

We gratefully thank Dr. Lisa Y. Armitige for providing all reference *M. tuberculosis* strains used in this study.

## REFERENCES

1. World Health Organization. Global tuberculosis report 2018. Geneva, Switzerland.

2. Foster N, Vassall A, Cleary S, Cunnama L, Churchyard G, Sinanovic E. The economic burden of TB diagnosis and treatment in South Africa. Soc. Sci. Med. 2015; 130: 42–50.

3. Blumberg HM, Burman WJ, Chaisson RE, Daley CL, Etkind SC, Friedman LN, et al. American Thoracic Society/Centers for Disease Control and Prevention/Infectious Diseases Society of America: treatment of tuberculosis. Am. J. Respir. Crit. Care Med. 2003; 167: 603–662.

4. Velayati AA, Farnia P, Masjedi MZ. The totally drug resistant tuberculosis (TDR-TB). Int. J. Clin. Exp. Med. 2013; 6: 307–309.

5. Klopper M, Warren RM, Hayes C, Gey van Pittius NC, Streicher EM, Müller B, et al. Emergence and spread of extensively and totally drug-resistant tuberculosis, South Africa. Emerg. Infect. Dis. 2013; 19: 449–455.

6. Gygli SM, Borrell S, Trauner A, Gagneux S. Antimicrobial resistance in *Mycobacterium tuberculosis*: mechanistic and evolutionary perspectives. FEMS Microbiol. Rev. 2017; 41: 354–373.

7. Gröschel MI, Walker TM, van der Werf TS, Lange C, Niemann S, Merker M. Pathogen-based precision medicine for drug-resistant tuberculosis. PLoS Pathog. 2018; 14: e1007297.

8. CRyPTIC Consortium and the 100,000 Genomes Project. Prediction of susceptibility to first-line tuberculosis drugs by DNA sequencing. N. Engl. J. Med. 2018; 379: 1403–1415.

9. Kouchaki S, Yang Y, Walker TM, Walker AS, Wilson DJ, Peto TEA, et al. Application of machine learning techniques to tuberculosis drug resistance analysis. Bioinformatics 2018; 1–7.

10. Niederweis M, Danilchanka O, Huff J, Hoffmann C, Engelhardt H. Mycobacterial outer membranes: in search of proteins. Trends Microbiol. 2010; 18: 109–116.

11. Glickman MS, Cox JS, Jacobs WR Jr. A novel mycolic acid cyclopropane synthetase is required for cording, persistence, and virulence of *Mycobacterium tuberculosis*. Mol. Cell. 2000; 5: 717–727.

12. Takayama K, Wang L, David HL. Effect of isoniazid on the in vivo mycolic acid synthesis, cell growth, and viability of *Mycobacterium tuberculosis*. Antimicrob. Agents Chemother. 1972; 2: 29–35.

13. Velayati AA, Farnia P, Ibrahim TA, Haroun RZ, Kuan HO, Ghanavi J, et al. Differences in cell wall thickness between resistant and non-resistant strains of *Mycobacterium tuberculosis*: Using transmission electron microscopy. Chemotherapy 2009; 55: 303–307.

14. Rocha-Ramírez LM, Estrada-Garcia I, Lopez-Marin LM, Segura-Salinas E, Mendez-Aragon P, Van Soolingen D, et al. *Mycobacterium tuberculosis* lipids regulate cytokines, TLR-2/4 and MHC class II expression in human macrophages. Tuberculosis 2008; 88: 212–220.

15. Barry CE 3^rd^. Interpreting cell wall ‘virulence factors’ of *Mycobacterium tuberculosis*. Trends Microbiol. 2001; 9: 237–241.

16. Mendelson M, Walters S, Smith I, Kaplan G. Strain-specific mycobacterial lipids and the stimulation of protective immunity to tuberculosis. Tuberculosis 2005; 85: 407–413.

17. Cox JS, Chen B, McNeil M, Jacobs Wr Jr. Complex lipid determines tissue-specific replication of *Mycobacterium tuberculosis* in mice. Nature 1999; 402: 79–83.

18. López B, Aguilar D, Orozco H, Burger M, Espitia C, Ritacco V, et al. A marked difference in pathogenesis and immune response induced by different *Mycobacterium tuberculosis* genotypes. Clin. Exp. Immunol. 2003; 133: 30–37.

19. Caws M, Thwaites G, Dunstan S, Hawn TR, Lan NT, Thuong NT, et al. The influence of host and bacterial genotype on the development of disseminated disease with *Mycobacterium tuberculosis*. PLoS Pathog. 2008; 4: e1000034.

20. Krishnan N, Malaga W, Constant P, Caws M, Tran TH, Salmons J, et al. *Mycobacterium tuberculosis* lineage influences innate immune response and virulence and is associated with distinct cell envelope lipid profiles. PLoS One 2011; 6: e23870.

21. Parwati I, van Crevel R, van Soolingen D. Possible underlying mechanisms for successful emergence of the *Mycobacterium tuberculosis* Beijing genotype strains. Lancet Infect. Dis. 2010; 10: 103–111.

22. Portevin D, Gagneux S, Comas I, Young D. Human macrophage responses to clinical isolates from the *Mycobacterium tuberculosis* complex discriminate between ancient and modern lineages. PLoS Pathog. 2011; 7: e1001307.

23. Ritchie ME, Phipson B, Wu D, Hu Y, Law CW, Shi W, et al. *limma* powers differential expression analyses for RNA-sequencing and microarray studies. Nucleic Acids Res. 2015; 43: e47.

24. Park CY, Krishnan A, Zhu Q, Wong AK, Lee YS, Troyanskaya OG. Tissue-aware data integration approach for the inference of pathway interactions in metazoan organisms. Bioinformatics 2015, 31: 1093–1101.

25. Mishra A, Akhtar S, Jagannath C, Khan A. Pattern recognition receptors and coordinated cellular pathways involved in tuberculosis immunopathogenesis: emerging concepts and perspectives. Mol. Immunol. 2017; 87: 240–248.

26. Cabanski M, Steinmüller M, Marsh LM, Surdziel E, Seeger W, Lohmeyer J. PKR regulates TLR2/TLR4-dependent signaling in murine alveolar macrophages. Am. J. Respir. Cell. Mol. Biol. 2008; 38: 26–31.

27. Cheung BKW, Lee DCW, Li JCB, Lau YL, Lau ASY. A role for double-stranded RNA-activated protein kinase PKR in *Mycobacterium*-induced cytokine expression. J. Immunol. 2005; 175: 7218–7225.

28. Mundhra S, Bryk R, Hawryluk N, Zhang T, Jiang X, Nathan CF. Evidence for dispensability of protein kinase R in host control of tuberculosis. Eur. J. Immunol. 2018; 48: 612–620.

29. Deb A, Haque SJ, Mogensen T, Silverman RH, Williams BRG. RNA-dependent protein kinase PKR is required for activation of NF-κB by INF-γ in a STAT1-independent pathway. J. Immunol. 2001; 166: 6170–6180.

30. Fallahi-Sichani M, Kirschner DE, Linderman JJ. NF-κB signaling dynamics play a key role in infection control in tuberculosis. Front. Physiol. 2012; 3:170.

31. Bai X, Feldman NE, Chmura K, Ovrutsky AR, Su W-L, Griffin L, et al. Inhibition of nuclear factor-kappa B activation decreases survival of *Mycobacterium tuberculosis* in human macrophages. PLoS One 2013; 8: e61925.

32. Manson AL, Cohen KA, Abeel T, Desjardins CA, Armstrong DT, Barry CE 3^rd^, et al. Genomic analysis of globally diverse *Mycobacterium tuberculosis* strains provides insights into the emergence and spread of multidrug resistance. Nat. Genet. 2017; 49: 395–402.

33. Cohen KA, Abeel T, Manson McGuire A, Desjardins CA, Munsamy V, Shea TP, et al. Evolution of extensively drug-resistant tuberculosis over four decades: Whole genome sequencing and dating analysis of *Mycobacterium tuberculosis* isolates from KwaZulu-Natal. PLoS Med. 2015; 12: e1001880.

34. Eldholm V, Monteserin J, Rieux A, Lopez B, Sobkowiak B, Ritacco V, Balloux F. Four decades of transmission of a multidrug-resistant *Mycobacterium tuberculosis* outbreak strain. Nat. Commun. 2015; 6: 7119.

35. Izu A, Cohen T, Degruttola V. Bayesian estimation of mixture models with prespecified elements to compare drug resistance in treatment-naïve and experienced tuberculosis cases. PLoS Comput. Biol. 2013; 9: e1002973.

36. Dookie N, Rambaran S, Padayatchi N, Mahomed S, Naidoo K. Evolution of drug resistance in *Mycobacterium tuberculosis*: a review on the molecular determinants of resistance and implications for personalized care. J. Antimicrob. Chemother. 2018; 73: 1138–1151.

37. Stagg HR, Lipman MC, McHugh TD, Jenkins HE. Isoniazid resistant tuberculosis – a cause for concern? Int. J. Tuberc. Lung Dis. 2017; 21: 129–139.

38. Unissa AN, Subbian S, Hanna LE, Selvakumar N. Overview on mechanisms of isoniazid action and resistance in *Mycobacterium tuberculosis*. Infect. Genet. Evol. 2016; 45: 474–492.

39. Palomino JC, Martin A. Drug resistance mechanisms in *Mycobacterium tuberculosis*. Antibiotics 2014; 3: 317–340.

40. Almeida Da Silva PE, Palomino JC. Molecular basis and mechanisms of drug resistance in *Mycobacterium tuberculosis*: classical and new drugs. J. Antimicrob. Chemother. 2011; 66: 1417–1430.

41. Sun G, Luo T, Yang C, Dong X, Li J, Zhu Y, et al. Dynamic population changes in *Mycobacterium tuberculosis* during acquisition and fixation of drug resistance in patients. J. Infect. Dis. 2012; 206: 1724–1733.

42. Hazbón MH, Brimacombe M, Bobadilla del Valle M, Cavatore M, Guerrero MI, Varma-Basil M, et al. Population genetics study of isoniazid resistance mutations and evolution of multidrug-resistant *Mycobacterium tuberculosis*. Antimicrob. Agents Chemother. 2006; 50: 2640–2649.

43. Ramaswamy SV, Reich R, Dou S-J, Jasperse L, Pan X, Wanger A, et al. Single nucleotide polymorphisms in genes associated with isoniazid resistance in *Mycobacterium tuberculosis*. Antimicrob. Agents Chemother. 2003; 47: 1241–1250.

44. Vilchèze C, Jacobs WR Jr. The mechanism of isoniazid killing: clarity through the scope of genetics. Annu. Rev. Microbiol. 2007; 61: 35–50.

45. Cohen T, Becetta MC, Murray MB. Isoniazid resistance and the future of drug-resistant tuberculosis. Microb. Drug Resist. 2004; 10: 280–285.

46. Seifert M, Catanzaro D, Catanzaro A, Rodwell TC. Genetic mutations associated with isoniazid resistance in *Mycobacterium tuberculosis*: a systematic review. PLoS ONE 2015; 10: e0119628.

47. Ford CB, Sha RR, Maeda MK, Gagneux S, Murray MB, Cohen T, et al. *Mycobacterium tuberculosis* mutation rate estimates from different lineages predict substantial differences in the emergence of drug-resistant tuberculosis. Nat. Genet. 2013; 45: 784–790.

48. Borrell S, Gagneux S. Infectiousness, reproductive fitness and evolution of drug-resistant *Mycobacterium tuberculosis*. Int. J. Tuberc. Lung Dis. 2009; 13: 1456–1466.

